# Cell-to-cell expression dispersion of B-cell surface proteins displays genetic variation among humans

**DOI:** 10.1101/792606

**Authors:** Gérard Triqueneaux, Claire Burny, Orsolya Symmons, Stéphane Janczarski, Henri Gruffat, Gaël Yvert

## Abstract

Variability in gene expression across a population of homogeneous cells is known to influence various biological processes. In model organisms, natural genetic variants were found that modify expression *dispersion* (variability at a fixed mean) but whether such effects exist in humans has not been fully demonstrated. Here, we analyzed single-cell expression of four proteins (CD23, CD55, CD63 and CD86) across cell lines derived from individuals of the Yoruba population. Using data from over 30 million cells, we found substantial inter-individual variation of dispersion. We demonstrate, via *de novo* cell line generation and subcloning experiments, that this variation exceeds the variation associated with cellular immortalization. By association mapping, we linked the expression dispersion of CD63 to the *rs971* SNP. Our results show that human DNA variants can have inherently-probabilistic effects on gene expression. Such subtle genetic effects may participate to phenotypic variation and disease predisposition.

## INTRODUCTION

Quantitative genetics is challenged by the fact that many genetic variants do not systematically affect phenotypic traits, but rather predispose to these traits. For example, disease-causing mutations can be found among healthy carriers. Over many individuals, although it is statistically significant, the effect of such variants is low because some individuals tolerate the mutation. Such cases of incomplete penetrance can result from hidden factors that compensate for the mutation via gene-gene or gene-environment interactions. In this case the probability of a given mutation to manifest itself is not 100%, *i.e.* it has a probabilistic effect, but only *apparently* because, although they are not known, compensatory deterministic factors exist. In addition, using model organisms and single-cell measurements, we and others have shown that genetic variants can also have effects that are *inherently* probabilistic: even in a controlled context, mutations can sometimes affect only some cells or individuals^1–7^. Consequently, we previously introduced the concept of single-cell Probabilistic Trait Loci (scPTL)^8^. This term is analogous to Quantitative Trail Loci (QTLs), and defines genetic variants that control single-cell traits in a non-deterministic fashion. In multi-cellular organisms, an scPTL that changes the statistical distribution of cellular traits in a tissue may affect the overall phenotype of an individual in a probabilistic manner, as discussed earlier^8^. It is possible that such loci contribute to disease predisposition, and their existence in humans has therefore been a recent topic of investigation.

A specific case of scPTL is defined by genetic loci that affect the cell-to-cell *variability* of a trait. For gene expression traits, it is known that differences in mean expression levels are usually accompanied by differences in variability^1,9,10^. Special care is therefore needed to distinguish genetic loci modulating variability *per se* from those that affect mean expression, and thereby variability. In model organisms, this distinction can be made by applying dedicated genetic crosses^4,11^. In humans, however, co-variation of mean and variability cannot be decoupled experimentally. Even if single-cell data can provide fine-scale descriptions of the genetic control of gene expression^12^, loci affecting variability are sometimes identified simply because of their effect on mean expression^13^.

The existence of human scPTL of gene expression variability *per se* is supported by the observations of Lu *et al.*^14^ who identified loci that affect gene expression variability in primary T-cells of specific subtypes. These loci affected variability in ways that were not explained by a modulation of the mean. This work was therefore essential in establishing that scPTL with probabilistic effects exist in humans under physiological conditions. However, the nature of the variability observed among primary T-cells is complex, because it possibly includes various sources of heterogeneities (see Discussion). For this reason, we sought here to investigate variability among cells of the same type and grown in a controlled environment, using cultured cell lines derived from different donors. Although less representative of the variability that is present *in vivo*, this approach offers better control on the cells’ subtypes and environment, which helps to determine strict probabilistic effects.

To this end, we utilized lymphoblastoid cell lines (LCLs) of the 1000 Genome Project Consortium^15^. These cell lines have been extensively genotyped and, being derived from B-cells, they express surface proteins that can easily be quantified on single cells via immunostaining and flow cytometry. Using this resource, together with freshly-generated LCLs and subcloning experiments, we found that the level of expression dispersion of four cell-surface proteins (the low-affinity receptor for IgE CD23, the Decay-Accelerating Factor CD55, the tetraspanin CD63 and the co-stimulator CD86) differs between human individuals, and we identified a *cis*-acting SNP that affects CD63 expression variability independently of the mean. Our results illustrate effects of genetic variants that are missed by classical studies because they take place non-deterministically and at the level of single cells.

## RESULTS

### Terminology

The present study describes differences in gene expression both between cells and between individuals. It is therefore important to clarify the meaning of several terms that are used hereafter. *Expression variability* will refer to differences in expression level of a protein between cells of the same genotype, same cell type and subtype, and which are extracted simultaneously from the same environment; given that expression variability often co-varies with mean expression, we also use the term *expression dispersion* of Sarkar *et al.*^13^ to refer to the amount of expression variability that is not explained by the mean; an *individual* will refer to a human person; *variation* will refer to differences of a given scalar value, for example a summary statistics of single-cell values, between individuals having different genotypes; *cell line* will refer to a population of immortalized cells of the same cell-type that can be propagated *in vitro* and which derives from a single donor individual; *clone* will refer to a cell line deriving from a single primary cell extracted from an individual; *sample* will refer to a population of cells that were cultured together in a single well and collected in a single tube for investigation.

### Quantifying expression dispersion using millions of lymphoblastoid cells

Comparing cell-to-cell expression dispersion across individuals of a cohort presents several difficulties. First, it requires acquisitions on single cells that are all of the same type (or subtype) and share a common environment. Second, this cell-type and environmental context must be the same when analysing all individuals of the cohort. Third, as for any trait, *inter*-individual variation of expression dispersion can only be interpreted if *intra*-individual variation is also estimated. Finally, a large number of cells must be analyzed to obtain robust estimates of dispersion. Lymphoblastoid cell lines represent a powerful resource to face these challenges. They have been invaluable for characterizing human genetic variation, and they were widely used to map quantitative trait loci (QTLs) of various molecular and cellular traits, including gene expression^16^. LCLs are derived from B-lymphocytes through immortalization by Epstein-Barr virus (EBV) infection. The advantage of lymphoblastoid cells is that they express many B-cell-specific cell-surface proteins, for which monoclonal antibodies have been developed and that can therefore easily be quantified on single cells. By using fluorescent-conjugated antibodies and flow-cytometry, as routinely done in immunological studies, it is possible to quantify the expression of an antigen in tens of thousands of cells in minutes. This can provide robust estimates of expression dispersion. Moreover, since many of these cell lines have been extensively genotyped, it is possible to search for an association between dispersion and genotype by applying linkage tests if differences in dispersion are observed between cell lines from different donors.

To make full use of this experimentally tractable system we selected cell lines from the Yoruba population in Nigeria that had previously been genotyped as part of the HapMap and 1000 Genomes Project^15^. We selected Yoruba samples, since previous studies had shown them to have the largest genetic diversity, which is favourable for genetic mapping. For all experiments, cells were cultured, fixed, immunolabelled with a fluorescent antibody directed against a cell-surface protein of interest, stained with DAPI and analyzed by flow-cytometry. We developed and applied a dedicated analysis pipeline to automatically gate cells of similar size and in the G1 phase of the cell-cycle, yielding distributions of single-cell protein expression levels (Fig. 1). The experiment, from culture to acquisition, was repeated several times so that the mean and coefficient of variation (CV) of these distributions could be compared within and between cell lines.

**Figure 1.**
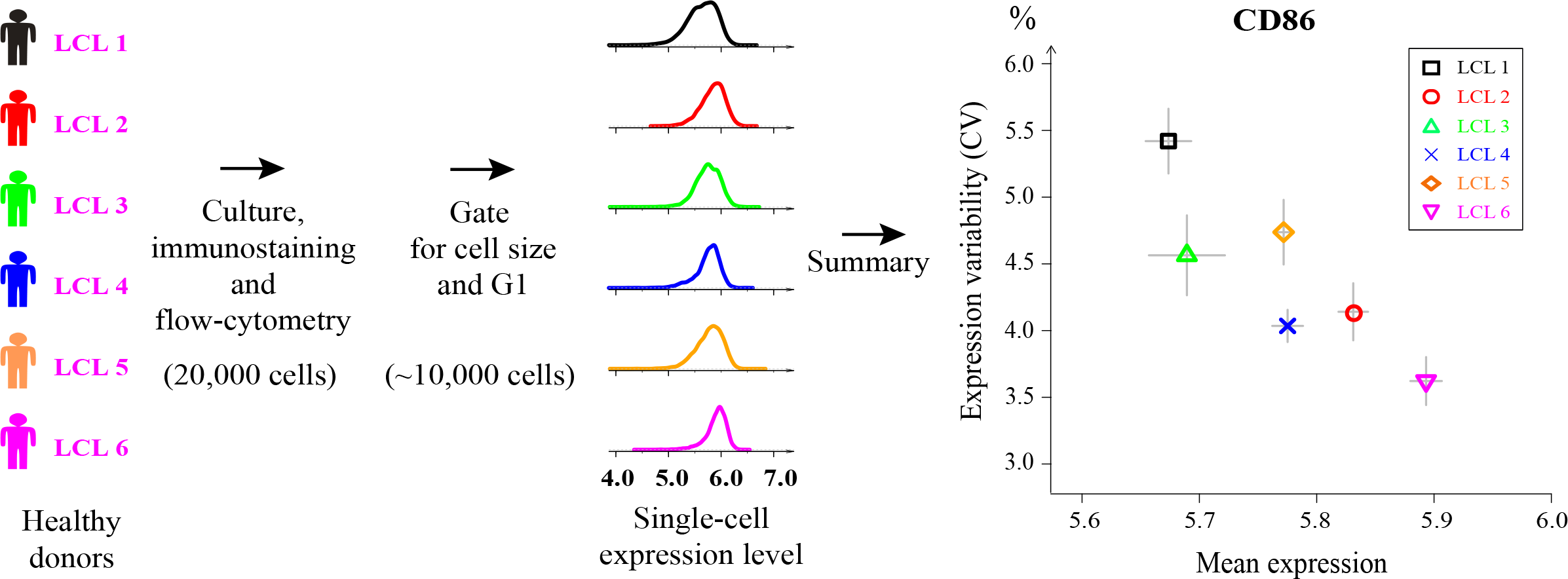
Experimental single-cell gene expression quantification in LCL lines. Cell lines from different healthy donors were cultured, fixed, immunostained with a fluorescent antibody, and analyzed by flow cytometry. A computational pipeline (see methods) automatically gated cells with similar size and in the G1 phase of the cell cycle, yielding distributions of expression values (log of fluorescence intensity) of the antigen protein of interest (here CD86). Summary statistics of the expression distributions were computed, such as the mean and coefficient of variation (CV = sd/mean). The experiment (culture, staining and acquisitions) was repeated several times independently to estimate intra-line variability (error bars: s.e.m., here *n* = 3 samples). Differences in CV between lines displaying similar mean levels of expression reflects different levels of cell-to-cell variability in expression (visible here for LCL4 versus LCL5).

### Expression variability of 18 surface proteins in cell lines from 6 unrelated individuals

As a first step, we started with a pilot survey using 6 cell lines from unrelated healthy Yoruba donors. We searched the literature and selected a set of 18 proteins meeting the following criteria: *i*) evidence of expression in LCLs, *ii*) availability of a validated monoclonal antibody suitable for fluorescent immunostaining and *iii*) these proteins participate to various biological functions of B cells, including disease-related processes. For each protein, we applied our labelling and analysis protocol and analyzed each cell line in triplicate (independent cultures and staining). We then computed the mean and CV of protein expression of the corresponding population of cells (Fig. 2A). Four proteins (CD2, CD9, CD37, CD79b) were not or very poorly detected in all 6 cell lines. Four proteins (CD40, CD46, CD59, CD80) were detected but displayed no marked differences in either expression mean or variability between cell lines. One protein (CD20) displayed mild variation in mean but not in variability. One protein (CD53) showed differences in variability between cell lines but it was weakly detected in all samples. Four proteins (CD5, CD22, CD38) displayed reliable differences of variability between cell lines, but this variation was fully correlated to variation of the mean (lower CV at higher mean). The remaining 5 proteins (CD19, CD23, CD55, CD63, CD86) were reliably detected, displayed variation in both mean and CV of expression, and, interestingly, variation in CV could not be directly attributed to variation of the mean. For example, the CV of CD23 expression decreased linearly with its mean expression across five cell lines but was significantly lower in the sixth cell line (blue color in Fig. 2); and for CD55, one cell line (orange color in Fig 2) displayed a larger CV but a similar mean expression as compared to two other cell lines. Example distributions are shown in Fig. 2B. Thus, our survey identified five proteins that displayed different levels of expression dispersion among six Yoruba cell lines.

**Figure 2.**
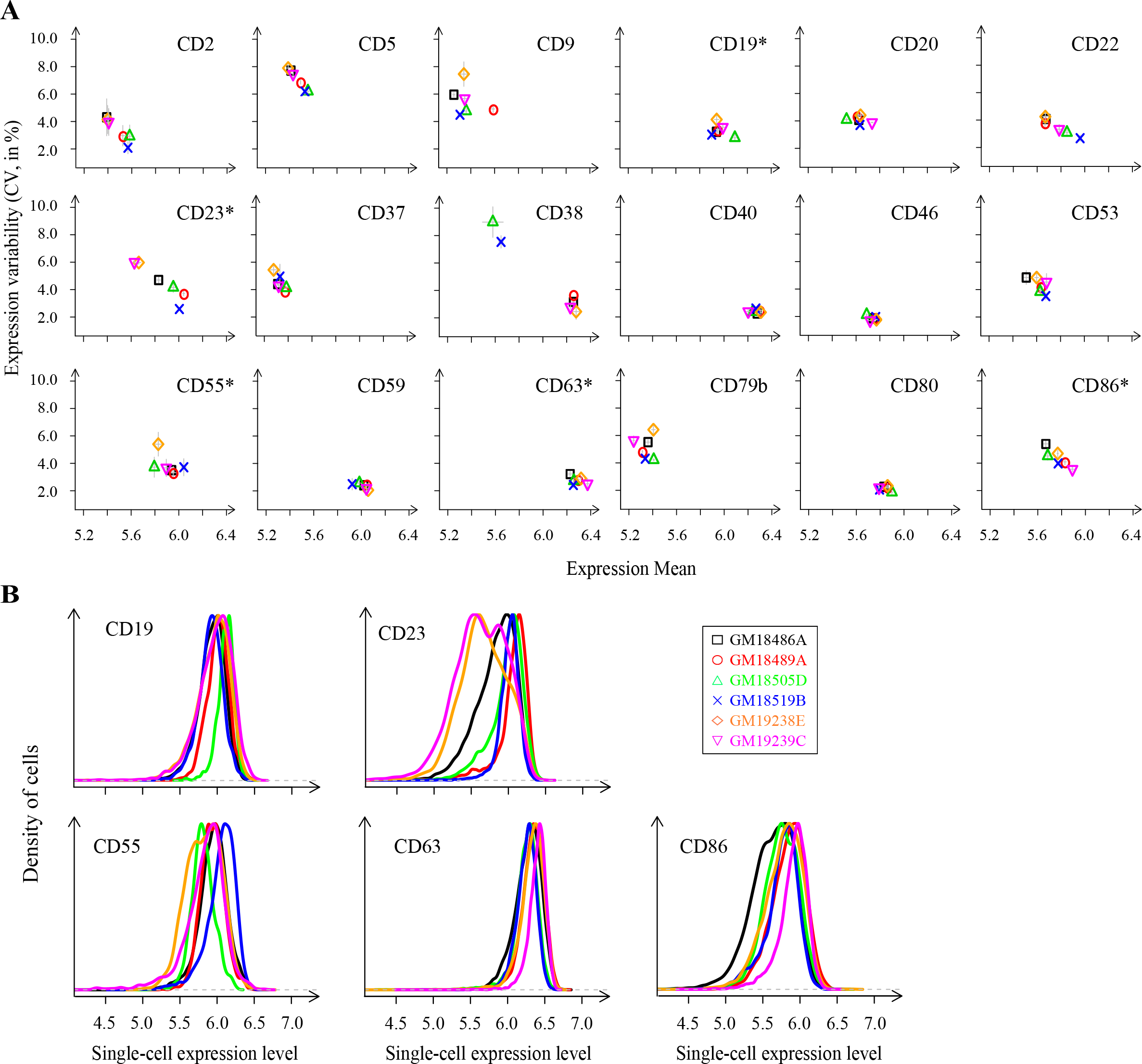
Single-cell expression of surface proteins in cell lines from unrelated individuals. **A**) Six cell lines from unrelated individuals (one per colored symbol) were immunostained for the indicated protein and analyzed by flow-cytometry to estimate CV and mean expression in cell populations. Grey error bars: s.e.m. between biological replicates of the same cell line, *n* >= 3 except for CD38 where n = 1 for one cell line and n = 2 for 4 cell lines). **B**) Distributions of single-cell expression values for 5 of the proteins shown in A. Each distribution corresponds to one randomly-chosen sample of the indicated protein and cell line (same color as in A).

### Single-cell expression distributions display specific patterns of variation

We decided to further characterize the single-cell expression properties of four of the five proteins: CD23, CD55, CD63 and CD86. We cultured, immunostained and analyzed a total of 50 Yoruba cell lines, in 6 biological replicates for each antigen and cell line. Note that, to avoid any biases of signal amplifications or crosstalk between fluorescent channels, each sample was stained with only one, directly-labelled antibody. We processed the data as above to derive distributions of expression values of G1-gated cells. We first plotted, for all samples, the CV (variability) of these distributions as a function of the mean (Fig. 3A). The four proteins clearly differed in their pattern of variation. CD23 was very poorly expressed in two cell lines and displayed wide variation of mean expression across the remaining 48 lines. Its CV of expression varied among moderately-expressing cell lines and decreased abruptly in cell lines having high mean expression. The spread of mean expression and CV was also large for CD55 and CD86, with an important distinction: for CD55, the CV decreased linearly with the mean, whereas for CD86, variation of CV and mean were independent (Fig. 3A). CD63 displayed the lowest variation in both mean and CV, although the CV was clearly elevated in three cell lines. We quantified expression dispersion of these three proteins by applying a locally-linear (lowess) regression to the CV vs. mean dependency. An example of this regression is shown in Fig. 3B. We used the residuals of the model (noted CV | mean, or *conditioned CV*) as estimates of expression dispersion. Figure 3C shows the spread of variation among cell lines of expression dispersion values. This variation was significantly larger than the variation observed among replicates of the same line (error bars). Of the four proteins, CD86 was the one displaying the largest line-to-line variation of expression dispersion.

**Figure 3.**
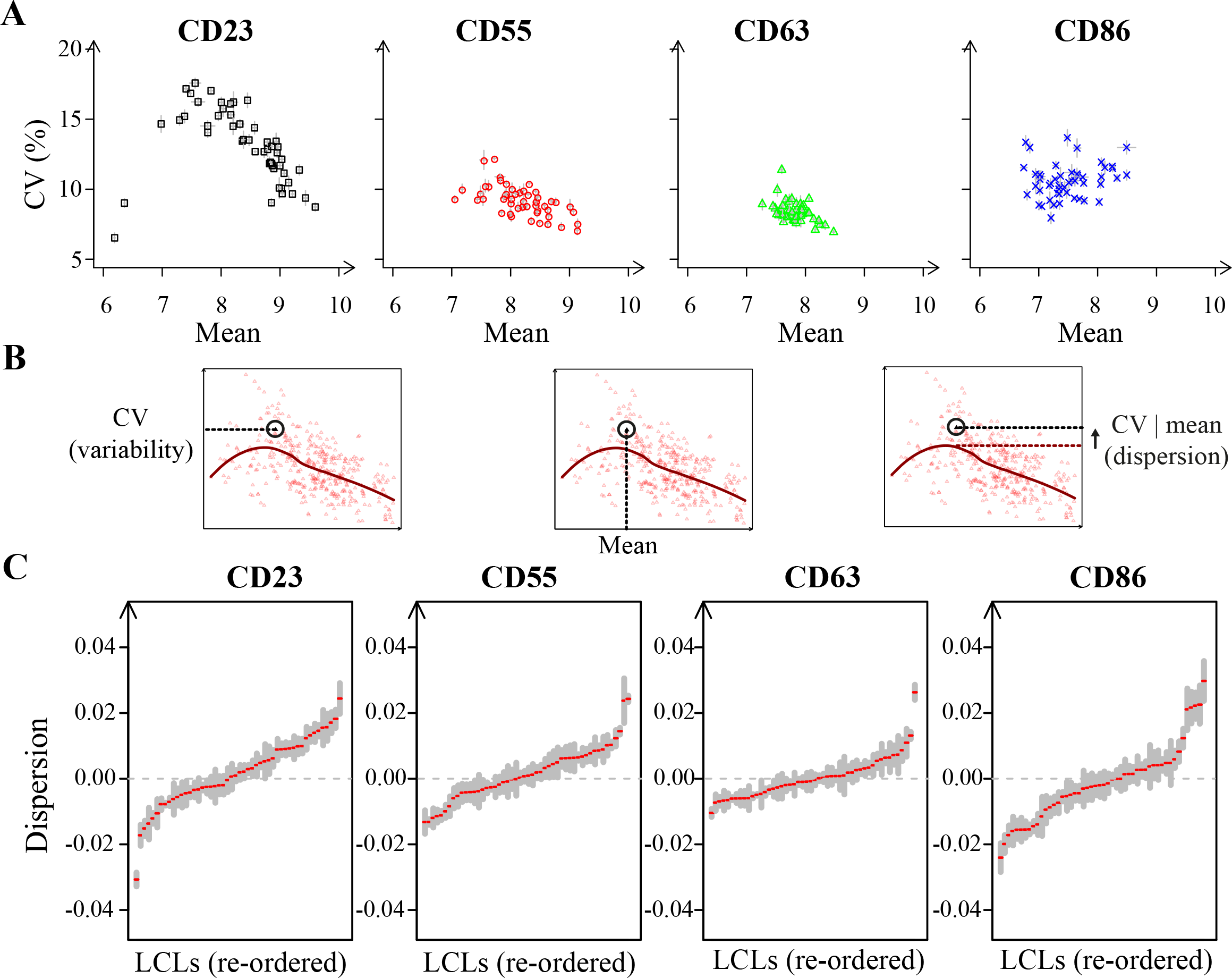
Patterns of variation in mean expression and CV across 50 individuals. **A)**For each sample (at least 6 per antibody and cell line), the CV and mean expression were computed. Each dot represents average values computed for each cell line and antibody combination (error bars = s.e.m). **B**) Lowess regression applied to the data (here for CD55 expression). The model was fitted to the entire dataset and for each sample, residuals were extracted (CV | mean = difference between the observed CV and the CV predicted by the model given the observed mean expression). **C**) Distributions of expression dispersion of the indicated proteins in 50 Yoruba cell lines. Each red tick corresponds to the average value of dispersion across biological replicates. Grey shaded bar = +/− s.e.m. (*n* >= 6).

### Expression dispersion of different proteins partially co-vary across individuals

We asked if variation of expression dispersion was correlated between proteins. For example, whether individuals with elevated dispersion for CD23 also display high dispersion for CD55. To address this, we computed all pairwise correlations between dispersion values in the 50 LCLs (Fig. 4). We found significant positive correlations for three pairs of proteins (CD23/CD63, CD23/CD86, CD55/CD86), and clearly no correlation for one pair (CD55, CD63). This reveals that co-variation between dispersion of different proteins exists, but only partially.

**Figure 4.**
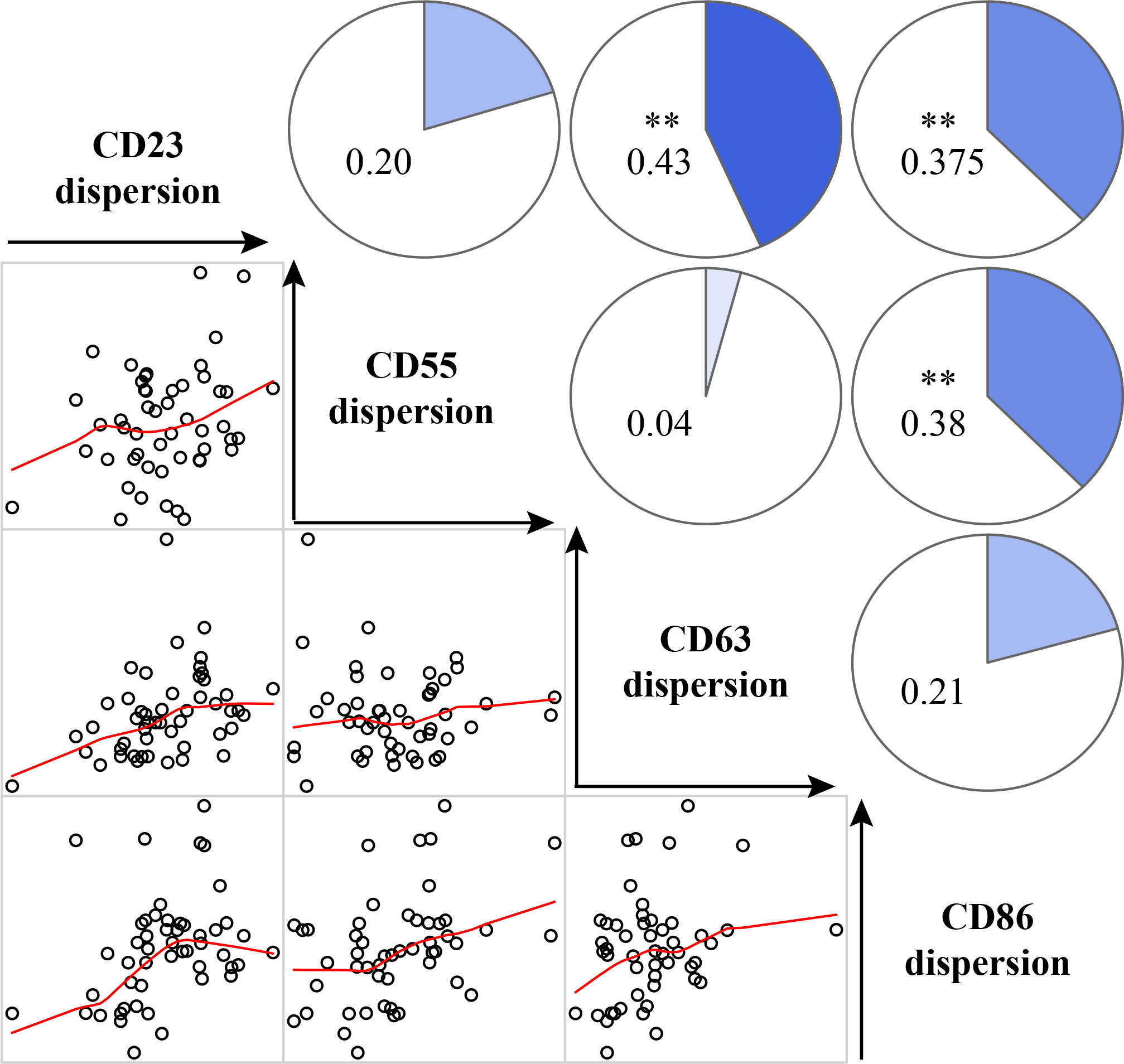
Correlation between expression dispersions of different proteins. Each dot plot represents expression dispersion values (CV | mean) of two proteins (which names appear on the diagonal) across 50 Yoruba LCLs (one per dot). Red line: smoothed regression. Pie charts and values correspond to Spearman rank correlation coefficients. Those labelled as (**) are significant at *p* < 0.01.

### Cell lines from some but not all Yoruba individuals contain both high- and low-expressing CD23 cells

We made the interesting observation that, in some but not all LCLs, the distribution of single-cell CD23 expression was bimodal (see example in Fig. 5A). In such cases, the mean and CV of the distribution do not capture all properties of single-cell expression. We therefore sought to provide a more comprehensive description by fitting a mixture model to the observed data. As illustrated in Fig. 5A, the model consisted of two Gaussians that each described a subpopulation of cells. The model was fully defined by five parameters: the means (*μ*_*1*_, *μ*_*2*_) and standard deviations (*σ*_*1*_, *σ*_*2*_) of the Gaussian components, and the proportion of cells belonging to the first component (*p*_*1*_). By fitting model parameters to the data of each CD23-stained sample, we compared how these distributions differed between cell lines. Hierarchical clustering of the cell lines based on parameter values revealed three groups of lines (Fig. 5B-C). Cluster 1 contained only three LCLs and was characterized by a majority of cells with low CD23 expression and a right tail of cells at higher CD23 levels. A second group containing half the LCLs had a mirrored pattern, with a bulk of cells at high expression and a left tail of low-expressing cells. For the remaining 22 LCLs, two subpopulations of cells defined by distinct modes of expression clearly co-existed. Thus, cell populations can differ not only in expression dispersion but also in expression bimodality, where two distinct states can be identified in some individuals but not in others. For CD23, which is a low-affinity receptor for IgE, bimodality implies that one cell type simultaneously generates highly- and poorly-responsive cells to low levels of IgE. Our results suggest that this duality can be pronounced in some but not in all individuals.

**Figure 5.**
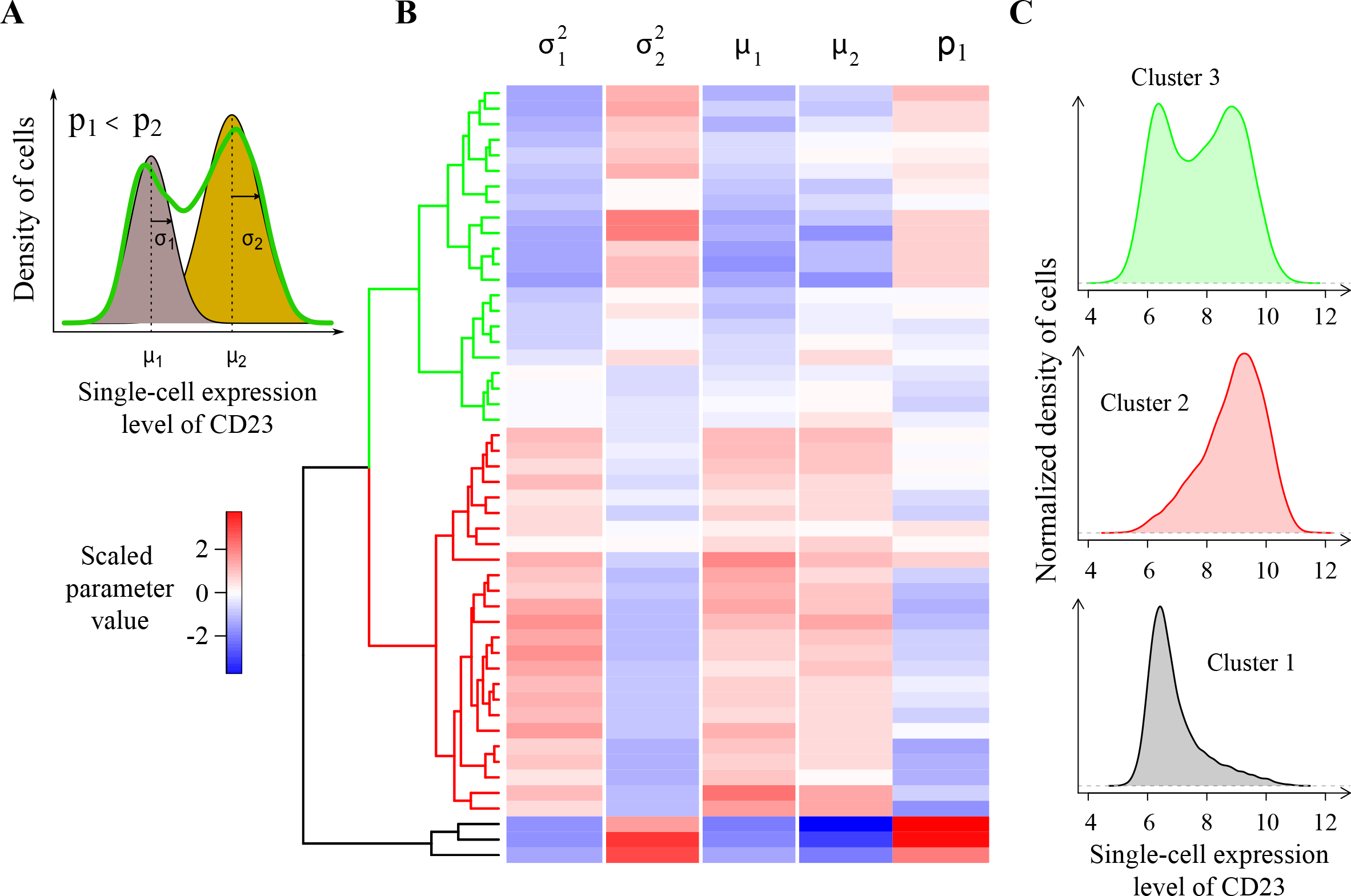
Variation in the bimodality of CD23 expression. **A)** Gaussian Mixture Model (GMM) describing the distribution of single-cell CD23 expression. Green curve: density curve of observed expression values in a sample from cell line GM18853B. Shaded distributions: Gaussian components of the model fitted to the data. **B**) Hierarchical clustering of LCLs based on GMM parameters. For each cell line, parameter values of replicate samples were averaged, centered to zero and scaled. Hierarchical clustering was applied using complete linkage. Note that colors are used to compare cell lines, they are not appropriate to compare parameters (due to scaling). **C**) Each distribution shows the observed data for a randomly-chosen sample belonging to one of the clusters (same color as tree shown in B). Clusters 1 to 3 contained 3, 25 and 22 LCLs, respectively.

### EBV-mediated immortalization does not explain inter-individual differences in expression dispersion or bimodality

The use of LCLs has been criticized because immortalization by EBV modifies cellular regulations as compared to non-infected primary cells. For example, CD23 is known to be upregulated in EBV-activated cells^17^. It is possible that EBV modifies not only the mean expression of host-cell genes but also their expression dispersion. In this case, independent immortalization events may generate cell lines with different levels of expression dispersion, regardless of original donor. It is therefore important to determine if the different levels of expression dispersion that we observed here could result from different outcomes of the immortalization process. To this end, we performed two complementary series of experiments.

We first reasoned that if differences observed between cell lines are significantly caused by inter-individual variation, then only few differences should be observed among cell lines originating from the same donor. Such variation between different cell lines from the same donor cannot be directly estimated from Yoruba LCLs because a single LCL is available per donor. We therefore generated additional cell lines from two unrelated and healthy donors. Primary cells from blood samples were infected with EBV and twelve independent cell lines were established from each donor. We cultured and processed these lines to quantify the expression of CD23, CD55, CD63 and CD86 at the single-cell level as above. To study variation within and between individuals, we first plotted the mean, variability (CV) and dispersion (CV | mean) of expression for CD55, CD63 and CD86 (Fig. 6A-B). For all three proteins, the two donors differed in both mean and CV values. Very importantly, the spread of variation between lines of the same donor was much lower than between lines of different donors. This was true not only for mean expression but also for CV and dispersion. In particular, all cell lines from one individual (donor 1 on Fig. 6) displayed markedly higher CD86 expression variability and dispersion than the cell lines from the other individual. Consistently, for all three proteins, variation in mean, CV and dispersion was larger among the Yoruba cell lines than among cell lines originating from a single donor. We analyzed CD23 single-cell expression distributions separately, taking into account bimodality. We fitted a Gaussian mixture model to the data of each sample and we analyzed variation of model parameters (Fig. 6C). For all five parameters of the model, variation among the Yoruba LCLs was significantly larger than the variation observed among lines of a single donor. These observations made on *de novo* LCLs demonstrate that, although intra-individual line-to-line variation in expression dispersion exists, it is lower than inter-individual variation.

**Figure 6.**
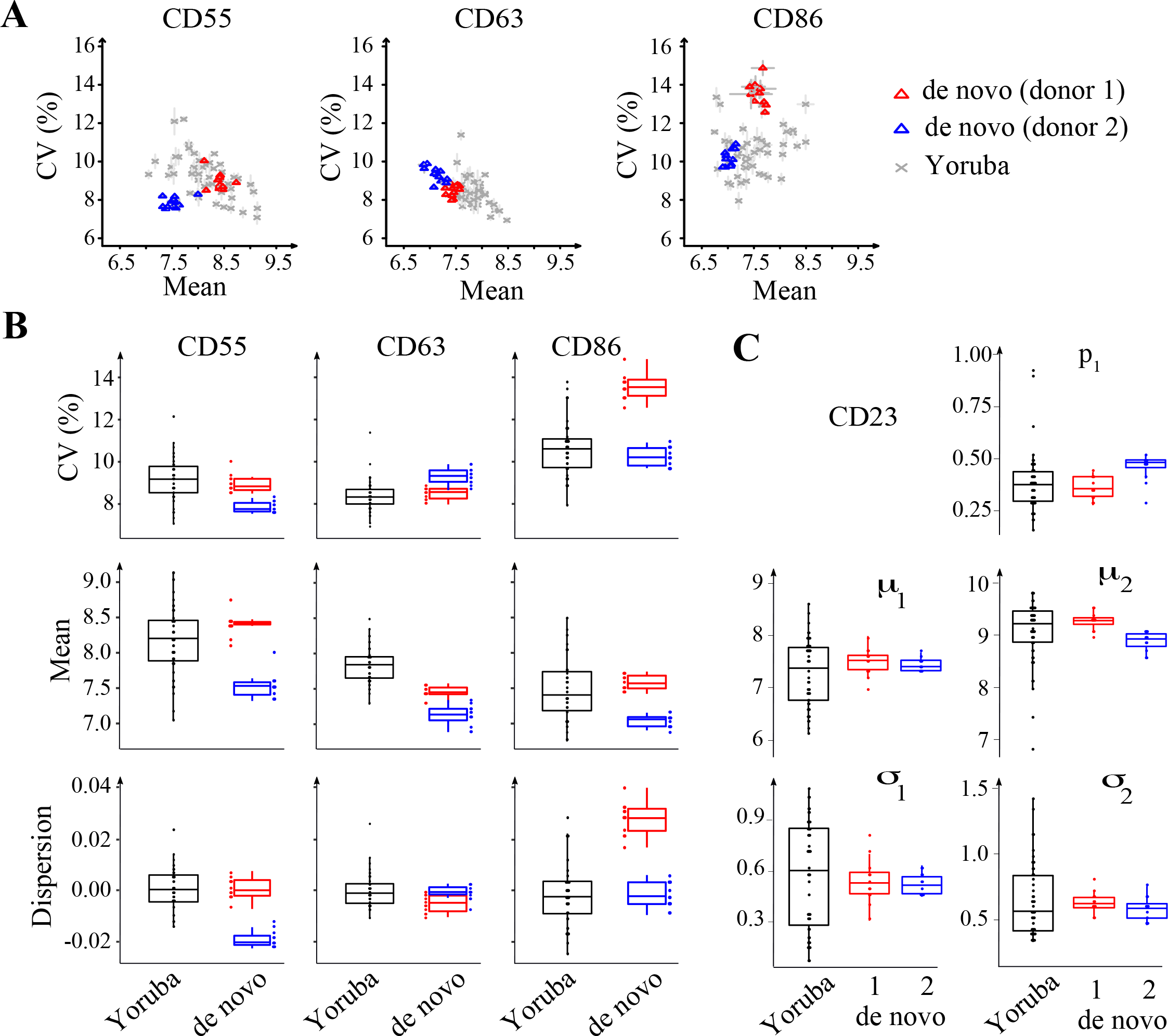
Intra- versus inter-individual variation of expression dispersion. Multiple LCLs were generated *de novo* using blood samples from two unrelated donors (blue and red), and expression mean and variability was compared to those observed in Yoruba LCLs. **A)** Dot plots of CV vs. mean expression of the indicated proteins. Each dot represents one cell line. Bars: s.e.m. (*n* = 2 independent cultures of each *de novo* LCL). **B**) Boxplot of CV, mean, and dispersion. Dispersion values correspond to CV|mean residuals computed from a lowess regression fitted to all samples. **C**) Boxplot of GMM parameter values fitted to distributions of single-cell CD23 expression (as in Fig. 5). Each dot represents one cell line. Values of replicate cultures were averaged. xxx define boxplot elements, e.g. center line, median; box limits, upper and lower quartiles; whiskers, 1.5x interquartile range; points, outliers xxx

In a second series of experiments we tested if elevated dispersion or pronounced bimodality in some Yoruba cell lines could potentially emerge as a consequence of multiple EBV-mediated immortalization events, since this process can result in polyclonal cell lines if more than one primary cell is infected and expands in a given sample. If different clones contained in a cell line expressed a protein at slightly different levels, this inter-clone variability would be seen as cell-to-cell variability at the level of the whole cell line. We therefore sought to i) determine, for some of the Yoruba cell lines, if they were monoclonal or polyclonal and ii) quantify expression dispersion in the context of monoclonality.

To distinguish between mono- and polyclonality, we took advantage of the somatic VDJ rearrangements and editing that take place during B-cell maturation. This process occurs exclusively *in vivo*, during a complex interplay between cell types within germinal centers, and generates the immunoglobulin specificity expressed by mature B-cells. Cell lines of the 1000 Genome Project were established by infecting blood cells with EBV in culture dishes *ex vivo*. Thus, any specific VDJ sequence is a signature of one cell that matured *in vivo* before cell line generation, and clonality of LCLs can be assessed by their genetic homogeneity at VDJ junctions: if the population is monoclonal only a single signature will be observed, while in a multiclonal population multiple signatures will be present. We adapted previously described protocols^18,19^ to simultaneously amplify several fragments of the VDJ junctions (Fig. 7A). A secondary amplification was then used to tag amplicons with indexing primers informing on the sample of origin and allowing for multiplexed 150bp paired-end Illumina sequencing.

**Figure 7.**
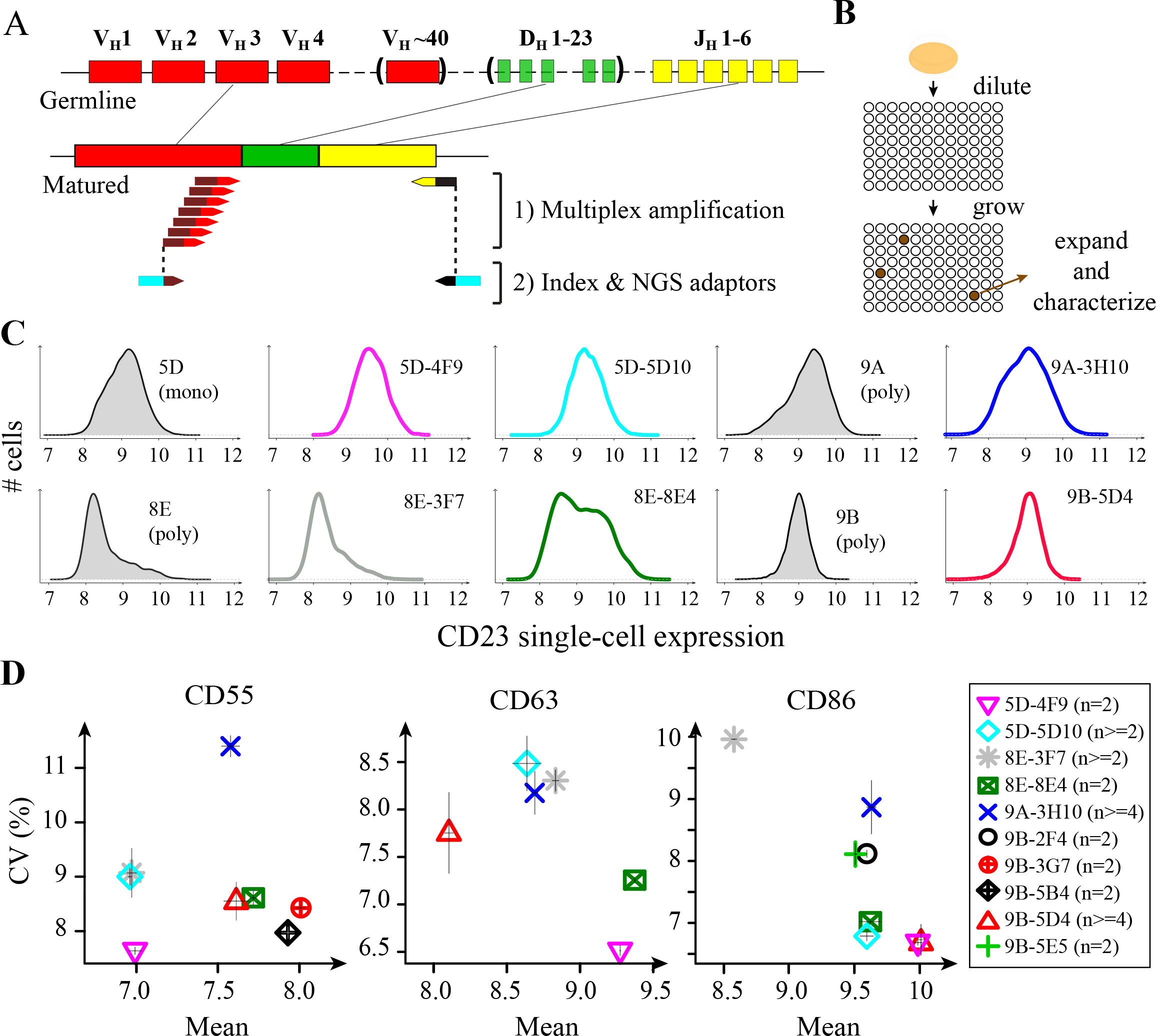
Subclones of LCLs also display variation in expression dispersion. **A)** Scheme of the clonality test applied to cell lines and subclones (see main Text and Supplementary Table S1 for results). **B)** Subcloning procedure. **C**) Single-cell expression distributions of CD23 in cell lines (shaded grey) and subclones (colored lines). 5D = GM18505D; 9A = GM18489A; 8E = GM19238E; 9B = GM18519B. **D**) Expression mean and CV of three proteins in ten monoclonal subclones. Bars: +/− s.e.m. *n*, number of independent samples. In C-D, subclones nomenclature indicates their origin (*e.g.* 5D-4F9 derived from 5D).

To study the relationship between expression dispersion and clonality in LCLs, we subcloned six cell lines by limiting dilution (Fig. 7B, see methods), which yielded a total of 27 subclones. Using the method described above, we genotyped these - presumably monoclonal - subclones and their parental lines, in duplicates, generating a total of 12,022,926 pairs of reads (median coverage per sample: 73,198 read pairs). To analyze this data, we developed a dedicated pipeline based on IgBlast^20^ that produced unique peptide sequences of the variable CDR3 region generated by VDJ rearrangements and editing (see methods). Results of this genotyping are shown in Supplementary Table S1. We found that two of the parental cell lines (GM18505D and GM18486A) were monoclonal: their genotype was homogeneous and was found in all of their subclone progenies. We also found that three Yoruba cell lines (GM18519B, GM18489A and GM19238E) were unambiguously polyclonal: their genotype was heterogeneous and genotypes of their progenies differed. Results on the sixth cell line (GM19239C) were inconclusive. Thus, the 1000 Genomes Project resource contains both monoclonal and polyclonal LCLs. This information may be important when interpreting variation in molecular traits among cell lines, especially for traits that are related to epigenetics. For example cell line GM19238, which we found to be polyclonal, was previously used in at least three studies of inter-individual variation of chromatin marks^21–23^. If these marks diverged between clones contained in the cell line, then the corresponding epigenomic trait values could be affected by the relative abundance of each clone in a sample.

Of the 27 subclones, 21 had a single CDR3 genotype and only 2 produced a clear mixture of genotypes (Supplementary Table S1). This validated the efficiency of the subcloning procedure. We therefore quantified expression dispersion in some of the subclones and their parental lines by immunostaining and flow cytometry as above.

Results for CD23 are shown in Figure 7C. Of the four parental cell lines that we re-analyzed, three showed high expression dispersion. Two of these lines were polyclonal (8E and 9A) and one was monoclonal (5D). The remaining cell line displayed very little CD23 expression dispersion and was polyclonal (9B). Thus, polyclonality does not correlate with CD23 expression dispersion in these Yoruba cell lines. We also analyzed six subclones. Five showed CD23 expression distributions that were fully consistent with the distributions observed on their respective parental lines. Importantly, the pronounced dispersion of polyclonal lines 9A and 8E was reproduced in their monoclonal derivatives 3H10 and 3F7. This excludes the possibility that expression dispersion in these two Yoruba LCLs was due to their polyclonality. In addition, subclone 8E4 displayed marked bimodality of CD23 expression (co-existence of low and high-expressing cells) despite its confirmed clonality. Thus, bimodality persists after single-cell subcloning and, in this example, it does not result from polyclonality.

As shown in Figure 7D, results for CD55, CD63 and CD86 also excluded a systematic association between polyclonality and expression dispersion. For all three proteins, we could find a pair of subclones that differed from one another regarding variability but not mean expression (3H10 > 5D4 for CD55, 8E4 > 4F9 for CD63, 3H10 > 5D10 for CD86). This demonstrates that LCLs can differ in expression dispersion of CD55, CD63 and CD86 despite being monoclonal.

Altogether, these observations on *de novo* and on monoclonal LCLs exclude the possibility that differences in expression dispersion between Yoruba LCLs simply result from variable outcomes of EBV-based immortalization. This motivated us to search for genetic factors that might underly these differences.

### Genetic mapping of expression variability and dispersion

We searched for association (genetic linkage) between DNA variants and gene expression dispersion using the available genotypes of the Yoruba LCLs. Of the fifty cell lines analyzed here, forty had been genotyped with phased and high-coverage data, eight were at an earlier stage with unphased data, and two were not characterized. We therefore applied linkage using either a high-quality map of SNPs covering 40 individuals, or a lower-density map covering 48 individuals. In each case, we searched for association between SNPs located in the vicinity of the gene of interest (+/− 2Mb from TSS) and expression traits: mean, variability, dispersion and, in the case of CD23, five fitted parameters describing bimodal distributions.

Results are summarized in Table 1. We found no association for mean expression levels but we did find associations for expression variability and dispersion. For CD23, dispersion was associated with 8 clustered SNPs but this association was marginally significant and supported by 4 individuals displaying reduced dispersion as compared to others (two individuals had barely-detectable expression, Supplementary Fig. S1A). For CD86, we found a significant association for both variability and dispersion using the 40 densely-genotyped individuals. This association corresponded to four heterozygous individuals displaying high dispersion. Notably, two individuals also displayed high dispersion but were not covered by the genetic map. To add them in the analysis, we genotyped them at the associated SNPs by PCR and sequencing. This revealed that they were homozygous, therefore eliminating the genetic association (Supplementary Fig. S1B). Finally, we found a significant association between CD63 variability and SNP *rs971* (Fig. 8A). This linkage was supported by both homozygous and heterozygous individuals, with one homozygous individual displaying high expression variability. Importantly, association was not accompanied by mean effect, and the genotypic groups also differed in expression dispersion (Fig. 8B). The mechanism by which *rs971* affects CD63 expression dispersion remains to be found. The SNP resides ~1.5Mb away from CD63, in the 3’UTR of SMUG1, a gene involved in base excision DNA repair (Fig. 8C). We inspected annotated positions of enhancers and transcription factor binding sites and found none overlapping *rs971*. Interestingly, according to dbSNP (www.ncbi.nlm.nih.gov/snp/) the *rs971* allele associated with high variability is not restricted to Yoruba but is present in all described human populations, with a minor allele frequency of at least 19%.

**Table 1:** Results of genetic association tests.

| Protein | Trait | #LCLs | Chromosome | Position (i) | SNPs | *p*-value | Conclusion |
| --- | --- | --- | --- | --- | --- | --- | --- |
| CD23 | CV\|mean | 40 | chr19 | 6246936 to 6277226 | rs75228027,<br>rs191354398,<br>rs116785610,<br>rs115321934,<br>rs116565827,<br>rs201284176,<br>rs114426250,<br>rs115581057 | 0.043 | marginal association (ii) (Supplementary Fig. S1A) |
| CD86 | CV | 40 | chr3 | 123464070 to 123492460 | rs116471535,<br>rs145139961,<br>rs150277702,<br>rs144503460,<br>rs116353991 | 0.0196 | Not validated with 2 more individuals (Supplementary Fig. S1B) |
|  | CV\|mean | 40 | chr3 | 123464070 to 123492460 | rs116471535,<br>rs145139961,<br>rs150277702,<br>rs144503460,<br>rs116353991 | 0.009 |  |
| CD63 | CV | 48 | chr12 | 54575458 | rs971 | 0.014 | QTL of variability (Fig. 8) |

(i): These SNP positions were retrieved from genotype data and may therefore differ from more recent physical maps.
(ii): *p*-value was corrected for the number of SNPs tested, but not for the number of traits investigated (8 for CD23).

**Figure 8.**
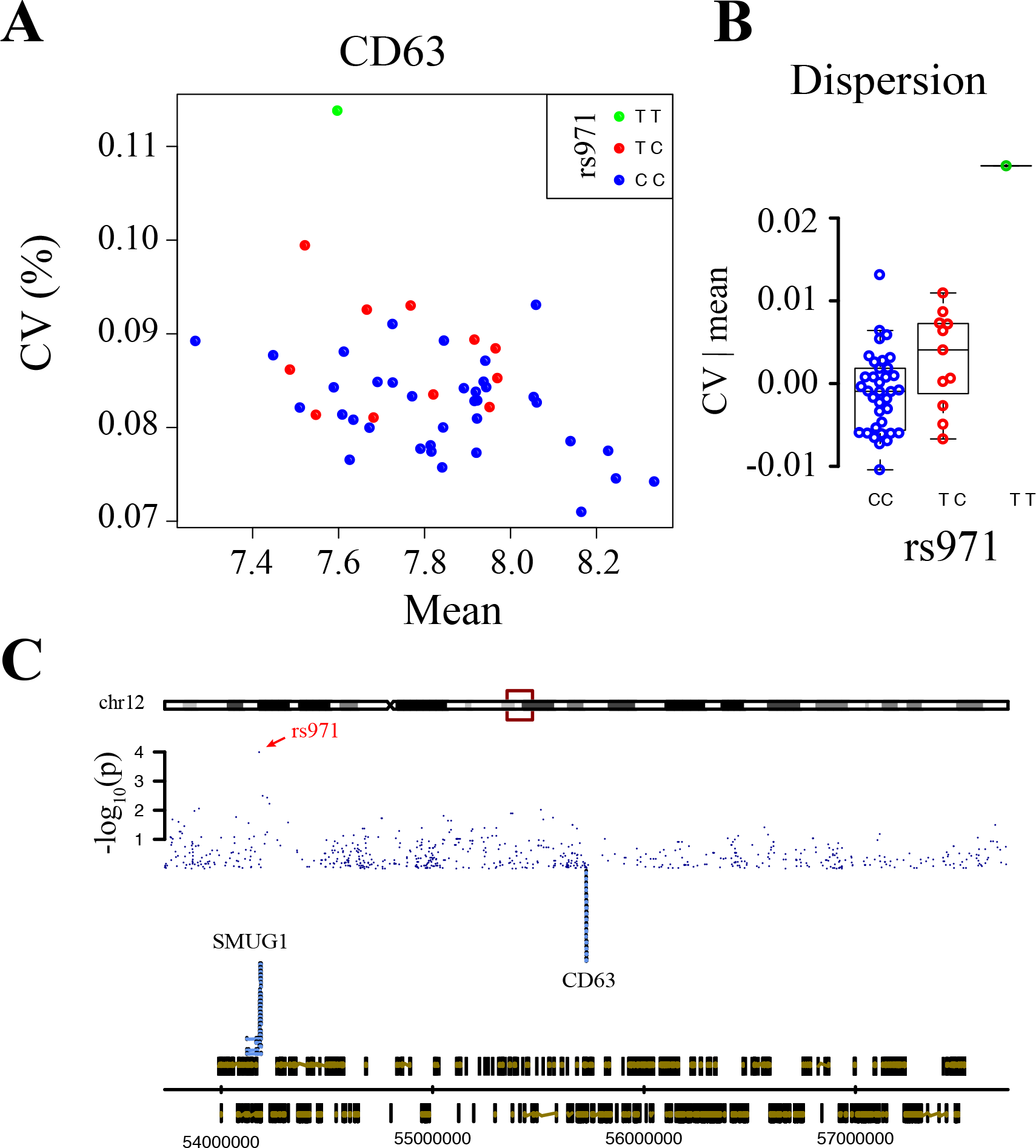
Genetic mapping of CD63 expression variability. **A**) Dot plot of CV vs. mean expression of CD63 in 48 LCLs, colored according to their genotype at SNP rs971. **B**) CD63 expression dispersion according to rs971 genotype. **C**) Genomic view of the locus. Blue dots, nominal linkage scores for association with CD63 expression variability (CV). Bottom track: genomic coordinates and genes positions. Middle track: transcripts of CD63 and SMUG1. Retrieved from Ensembl on 2019-06-11 using GenomeGraphs^46^.

## DISCUSSION

By quantifying a handful of cell-surface proteins in millions of individual lymphoblastoid cells, we found that the level of cell-to-cell expression variability can differ between healthy humans. For proteins CD23, CD55, CD63 and CD86, variation in mean expression did not explain the differences observed on variability levels, demonstrating variation of expression *dispersion* among humans. Expression of CD23 was bimodal for some individuals (co-existence of two subpopulation of cells with different mean expression) but not for other individuals. Analysis of *de novo*-generated cell lines and subcloning experiments excluded the possibility that expression dispersion and bimodality systematically resulted from variable outcomes of the immortalization process. Consistently, we found a *cis*-acting SNP linked to CD63 expression variability independently of the mean.

Our results complement the study of Lu *et al.*^14^ who analyzed single-cell expression of 14 proteins in various primary T-cell subtypes from unrelated donors. Their experimental design and ours have several features in common: antibody staining and flow-cytometry, a focus on dispersion rather than variability - although the term used by Lu *et al.* was *cell-cell expression variation (CEV)* - and coverage of a cohort of healthy individuals allowing genetic linkage analysis. The two studies also have important differences. Lu *et al.* used primary T-cells while we used lymphoblastoid cell lines; they quantified all proteins simultaneously by multicolor staining and compensation while we used single labelling; they interrogated ~800 SNPs that had previously been linked to immunological traits and that were located away from the gene for which expression was considered (*trans*-acting SNPs), while we scanned thousands of SNPs located at the gene locus (*cis*-acting SNPs); and their cohort included individuals from different origins. These specificities have implications on the interpretation of results. First, cells were not only from different types (T versus B) but were also extracted from a totally different context (primary versus cultured lines). In the case of Lu *et al.*, the data from primary T-cells directly reflects *in vivo* cell states and their work demonstrated that both genotype and age affect expression dispersion in a physiologically-relevant context. However, the nature of this dispersion is not easy to define in this case. Primary cells belong to a complex classification of subtypes and the definition of cell-cell variability heavily relies on how each subtype is defined. Lu *et al.* provided detailed reports by carefully stratifying the data (manual gating). Nonetheless, if a subtype is in fact structured in two or more subpopulations of cells, the variability that is measured can then include both stochastic variability within each subpopulation and differences in mean expression between subpopulations. In contrast, culturing cell lines *ex-vivo* avoids this difficulty because it generates cells that directly come from a single subtype, especially after subcloning as we did here. Our results therefore directly demonstrate the variation of dispersion for a strictly-defined subtype. Secondly, primary cells of different individuals obviously come from different environments (different human bodies). Lu *et al.* accounted for this by studying the same individuals at consecutive time points, and by using sex and age as covariates, revealing cases where age impacted dispersion. However, various other physiological differences may contribute to dispersion, such as diet, lifestyle, microbiome and the ‘exposome’. The effects of all such factors are excluded from *ex-vivo* cultured cell lines because the environment is artificially controlled over a prolonged time. Our results show that variation of dispersion is present despite this controlled context. Finally, unlike in primary cells, gene expression dispersion in LCLs may be affected by EBV infection, for example if EBV changes the amplitude or dynamics of expression in host cells over time. Such effects may shift dispersion levels as compared to non-infected cells and, accordingly, we observed substantial differences in dispersion among LCLs originating from the same donor. However, these differences were weaker than the ones observed between LCLs of unrelated donors. This shows that the possible contribution of EBV on expression dispersion does not dominate the contribution of donor-specific factors, such as genetic variants. The two studies are further complementary by illustrating the contribution of *trans*-acting SNPs in T-cells (Lu *et al.*) and a *cis*-acting SNP in B-derived cells (this work).

Recently, Sarkar *et al.*^13^ searched for *cis*-acting SNPs of expression variability in another cell type: induced pluripotent stem cells, which were derived from the Yoruba LCL lines analyzed here. The authors used single-cell RNAseq, generating data on many more genes but much fewer cells as compared to flow-cytometry. This study failed to identify SNPs affecting dispersion: although SNPs could be associated with variability, this association was fully explained by their effect on mean expression. We note another layer of complexity in this work: cellular reprogramming may generate variability and it would be interesting to compare various iPSC lines originating from the same LCL.

We previously described a statistical method (ptlmapper) that can use the full distribution of single-cell traits to specifically identify scPTL^7^. This method looks for scPTL that may affect any property of the single-cell expression distribution, and not only specific pre-defined features such as mean, variability or dispersion. We note that when we applied this method to the dataset reported here, we did not detect any statistically-significant association (not shown). This is not particularly surprising given previous tests on yeast data, where the method was complementary but not necessarily more powerful than applying QTL mapping on dispersion traits^7^.

What is the biological implication of the genetic modulation of CD23, CD55, CD63 and CD86 expression dispersion? Although the inter-individual differences in dispersion that we report here are subtle they may be important for two reasons. First, a slight probabilistic effect is negligible over small numbers, but becomes clearly significant over millions of ‘trials’. Given the lifetime of a human individual, its high number of B-cells and the known activity and reactivity of these cells, the number of times surface proteins are used - the number of ‘trials’ - is enormous. The second reason is that, for immune cells, what happens to one can matter to all because each cell can trigger a response that is collectively amplified. However, a possible whole-organism significance of our observations remains to be determined because i) genetic effects are known to be context-dependent and Petri dishes are radically-different environments than human tissues; ii) the subtle effects that we described here *ex vivo* may be hidden by variability resulting from the dynamics of the blood environment (cross-talks with other cell types, variation in nutriments, flow mechanics…); iii) since molecular and cellular networks can buffer fluctuations, dispersion at the level of one macromolecule does not necessarily imply phenotypic alterations.

The four proteins analyzed here have multiple functions. CD23 is the low-affinity receptor for IgE but can also bind CD21, major histocompatibility complex class II proteins and integrins; it has several isoforms and it can be cleaved to release a functional soluble form^24^. Its expression is known to be strongly activated by EBV-mediated activation^17^. Our observation that CD23 expression is bimodal in EBV-immortalized cell lines from some but not all individuals suggests that this strong activation might be heterogeneous in a genotype-dependent manner. CD55 is the Decay-Accelerating Factor, which protects cells from the complement system^25^. It has additional roles by transducing signals in T-cells^26^. CD86 is a co-stimulator of CD28-mediated T-cell activation^27^. CD63 is a tetraspanin with numerous regulatory roles^28^, including exosome-based co-stimulation of T cells. Thus, different degrees of CD55, CD86 or CD63 expression dispersion in B cells may in turn generate heterogeneities among T cells. Finally, genetic variation in expression dispersion may also have clinical implications. For example, CD55 and CD63 are helpful for the prognosis of various cancers^29–32^ but interpretation of these biomarkers may be complicated if their expression dispersion is substantial in some patients and not others despite a similar clinical situation. Furthermore, CD23 is the target of lumiliximab, a monoclonal antibody used to fight chronic lymphocytic leukemia (CLL)^33^. If the patient’s genotype modulates CD23 expression bimodality among CLL cells, as we observed here for LCL cell lines, then it probably also modulates the proportion of cells that are efficiently targeted by this treatment.

In conclusion, our observations are important because they demonstrate the existence of non-deterministic genetic effects on gene regulations in humans, and further work is needed to link such effects to physiological perturbations and disease.

## METHODS

### Lymphoblastoid Cell Lines

Yoruba LCLs were part of the HapMap/1000 Genomes panel and were obtained from the Coriell cell repository (Camden, NJ, USA). Cell lines references are: GM18486A; GM18487; GM18488B; GM18489A, GM18498B; GM18499B; GM18501C; GM18502B; GM18504B; GM18505D; GM18507E; GM18508E; GM18516B; GM18517D; GM18519B; GM18520B; GM18522C; GM18523C; GM18852C; GM18853B; GM18855C; GM18856D; GM18858C; GM18859C; GM18861C; GM18862C; GM18867B; GM18868B; GM18870B; GM18871B; GM18873B; GM18874B; GM18912C; GM18913B; GM18916C; GM18917B; GM18933B; GM18934B; GM19098B; GM19099B; GM19107B; GM19108B; GM19140B; GM19141C; GM19192B; GM19193C; GM19203C; GM19204B; GM19238E; GM19239C. They corresponded to healthy adult donors from 25 YRI families who were unrelated except GM18913B who was the child of GM19238E. Additional LCLs were generated from two individual donors as follows. Primary B cells were freshly isolated from two adult human blood buffy coats by density gradient centrifugation, followed by positive selection using anti-CD19 Immunomagnetic beads (STEMCELL Technologies). Purified B cells (2.10^5^ cells) were exposed to supernatants of HEK293_EBV.B95-8_ EBV-producing cells^34^ at a multiplicity of infection (MOI) of 0.5 infectious particles per B cell, overnight at 37°C and then washed once with PBS and cultured with RPMI supplemented with 20% FBS until establishment of LCLs (25 days). Twelve LCLs were established from each donor.

### Subcloning of LCL lines

We isolated monoclonal subclones from five of the LCLs (GM19238E, GM18486A, GM18489A, GM18519B, GM19239C) by counting and diluting cells to a concentration of approximately 1-10 cells/ml. 100 μl of this dilution were then added to the wells of 96-well plates that already contained 100 μl of conditioned medium. Conditioned medium was obtained by culturing LCLs to a concentration of 300,000-500,000 cells/ml, pelleting the cells by centrifugation, collecting the supernatant and filtering it through an 0.2μ Stericup filter (Merck-Millipore). Diluted cells were then left to grow in 200 μl 50% conditioned media for 3-4 weeks, until growth was observed in some wells, occasionally supplementing the wells with 50% conditioned media to replace evaporated liquid. From the wells where growth was observed, cells were re-plated into 6-well plates and subsequently 25ml flasks. Finally, vials from each subclone were frozen for further analysis.

### Clonality test by PCR and NGS sequencing of CDR3

We established a protocol for PCR-based amplification and multiplexed sequencing of the framework 3 VH region based on previous publications^18,19^. To this end, we isolated DNA from cell lines by transferring the cells to 1.5ml Eppendorf tubes, pelleting them by centrifugation, and lysing them overnight at 65°C in 500μl lysis buffer (100mM Tris pH=8.5, 5mM EDTA, 0.2% SDS, 200mM NaCl), complemented with final concentration of 0.1mg/ml ProteinaseK. The next day, DNA was precipitated with 350μl isopropanol at −20°C for 2h, centrifuged for 30min, the supernatant removed, and the pellet washed with 70% ethanol. After additional centrifugation and removal of supernatant the pellet was dried and resuspended in 200μl TE. We then amplified the region of interest from 1μl of DNA by a PCR reaction of 35 cycles in a final volume of 15μl containing an equimolar mixture of primers (2μM final concentration per primer) extended with Illumina adapters (see primer list in Supplementary Table S2), Platinum Taq (ThermoFisher), and 1.5 mM final MgCl2 concentration. We then performed PCR cleanup by incubating 5μl of the PCR reaction with 1μl Exonuclease I (NEB) for 30 minutes at 37°C, and inactivating the enzyme by a 15-minutes incubation at 80°C. We then used 3μl of cleaned PCR product as template for a second round of PCR using Illumina Nextera XT Index primers in 50μl volume. Amplicons quality was checked on agarose gel and we pooled 35μl from each PCR and performed Exonuclease I cleanup as above. The samples were then split into 4 tubes, run through a Macherey-Nagel PCR cleanup column and each sample was eluted in 30μl. All products were then pooled, diluted to approx. 150 ng/μl and sequenced by 150 bp paired-end sequencing on a MiSeq Illumina sequencer. Reads quality was assessed with FastQC (http://www.bioinformatics.babraham.ac.uk/projects/fastqc/, last accessed November 2015). Overlapping paired-end reads were merged using PEAR^35^. We then used bioawk (https://github.com/lh3/bioawk) to extract assembled reads with an average Phred score ≥ 33 and unassembled reads were discarded. FASTQ files were converted into FASTA files using a custom bash function. For mapping, we downloaded Fasta files from http://www.ncbi.nlm.nih.gov/igblast/showGermline.cgi in November 2015 (now migrated at http://www.imgt.org/) and we used standalone IgBLAST version 1.4. with commands C1-C4 of Supplementary Table S3 to create a database of V, D and J human segments and to assign each read to a specific gene segment, as defined in the IMGT ontology (http://www.imgt.org). Output files were then parsed using a parser of the maintained Galaxy toolshed https://testtoolshed.g2.bx.psu.edu/view/davidvanzessen/igblastparser_igg/176ce910f659 in order to obtain files having one line per read. We then filtered the data by keeping reads meeting all of the following conditions: “CDR3 Found How” ≠ “NOT_FOUND”, “VDJ Frame” ≠ “N/A” and “Top D Gene” ≠ “N/A”. We excluded singletons (CDR3 sequence covered only once). We removed reads for which CDR3 length was shorter than 5 amino-acids or for which the conserved CDR3/FR4 Phe/Trp-Gly-X-Gly motif in the J region was not present^36^. We considered that a unique CDR3 sequence corresponded to one clone^37^. The percentage of reads matching a given CDR3 sequence was used to define the representativity of the corresponding clone in the sample. If multiple CDR3 sequences reached 20% of representativity, the sample was considered polyclonal; if the major clone reached 80%, the sample was considered monoclonal; otherwise we considered the data to be inconclusive. In Supplementary Table S1, only sequences reaching 20% of representativity are shown.

### LCLs cultivation and immunostaining

Cells were cultured in Roswell Park Memorial Institute medium (RPMI 1640, GlutaMAX, Thermo Fisher Scientific) supplemented with 15% fetal calf serum (FCS, Eurobio), antibiotics (100 μg/ml penicillin, 100 μg/ml streptomycin). The cells were incubated at 37 °C in a humidified CO2 incubator (5% CO2) in 25 cm^2^ culture flasks (vented caps). Depending on cell growth, cell lines were diluted two to three times per week. Cell density was maintained between 300,000 cells/ml to less than a million cells/ml in 10 ml final volume of medium to optimize their growth. Although fixation is not mandatory for quantification, we noticed that it reduced day-to-day technical variability and we therefore applied it systematically. For fixation, cells were counted using KOVA slides after a 2-days growth phase (post-passage). Two million cells were collected, adjusted with PBS to a 10 ml volume, followed by centrifugation at 1,000 rpm for 5 min at room temperature. Cell pellets were washed with 4 ml PBS, and centrifuged as previously. They were then resuspended in 1.2 ml 4% Paraformaldehyde (PFA) in PBS and placed on a roller-tube agitator for 20 min at room temperature. Then 4 ml of PBS + 2% fetal calf-serum (FCS) were added and cell suspensions were centrifuged at 2000 rpm for 2 min. Pellets were resuspended in 2 ml of PBS + 2% FCS prior to immunolabelling. All labelling steps were performed in 96-well V-bottom plates in which 150 μl of fixed cells (approx. 150,000 cells) were deposited per well. Cells were centrifuged for 2 min at 2000 rpm, and resuspended in 100 μl of PBS + 10% FCS. Subsequently, each well was completed to 200 μl as follows: unlabelled cells (control) were supplemented with 100 μl of PBS + 10% FCS, while for labelling cells we added 100 μl of PBS + 10% FCS containing the antibody at appropriate dilution (See Supplementary Table S4). The plate was then incubated for 1 hour at 4°C followed by a 2000 rpm centrifugation for 2 min. Cells were washed with 200 μl of PBS, followed by another round of centrifugation. Finally, cells were resuspended in 230 μl of freshly diluted DAPI (2.5 μg/ml) in PBS + 1% FCS.

### Single-cell flow cytometry acquisitions

Plates were analysed on a BD FACSCanto II (BD Biosciences) flow cytometer equipped with a High Throughput Sampler (HTS). Acquisitions parameters were configured as follows: area scaling values of forward scatter (FSC) and side scatter (SSC) channels were optimized with photomultiplier tubes (PMT) voltages at 295 mV and 415 mV, respectively. Fluorescent PMT voltages were adapted for each antibody. DNA content was estimated from DAPI signal acquired in a Pacific Blue filter with PMT set at 280 mV. For each sample, 20,000 cells were acquired.

### Analysis of flow cytometry data: dataset structure

The study included three sets of experiments that were performed independently and that we treated separately. Set 1 covered the expression levels of 20 proteins (CD2, CD5, CD9, CD19, CD20, CD22, CD23, CD37, CD38, CD40, CD46, CD53, CD55, CD59, CD63, CD79b, CD80, CD86, IgKappa, ROR1) in 6 LCLs of the 1000Genome project (GM18519B, GM19238E, GM18489A, GM18486A, GM18505D, GM19239C). Set 2 covered the expression levels of 4 proteins only (CD23, CD55, CD63, CD86) in 50 LCLs of the 1000Genome project, in 24 LCLs originating from 2 ungenotyped donors, and in 10 subclones derived from Yoruba LCLs of the 1000 Genome Project. Note that for CD63 and CD86, this set contains heterogeneities between the series of 50 LCLs and the series of subclones: some cell lines were analyzed in both series and their mean values were shifted (Supplementary Fig. S2). It would therefore not be appropriate to compare single-cell distributions between the two series. Set 3 corresponded to quantification of CD23 in 6 subclones using a FITC-conjugated antibody (ref 561146 from Pharmingen, Supplementary Table S4) instead of the APC-conjugated anti-CD23 antibody (BD BioSciences, Supplementary Table S4). The following framework was separately applied to each set.

### Analysis of flow cytometry data: data filtering and normalization

All analysis was done using R (www.r-project.org) and packages flowCore^38^ and flowStats^39^ from Bioconductor (www.bioconductor.org). For a given experimental data set, data was pre-processed with the following steps. (i) Removal of events with saturated signals. This was done by setting, for each channel, a lower and an upper bound of signal intensity. These boundaries corresponded to the 5th and 95th percentile of all single-event values pooled from all samples. Events falling outside these boundaries in one or more channel were discarded. (ii) For channels where this procedure generated a negative lower bound, all values were augmented by adding to them the absolute value of this lower bound. This avoided negative values in downstream analysis. (iii) Removal of unstained DAPI events. For this, we identified the first motif of peak-valley-peak in the distribution of DAPI intensities by building a distribution density using the stats::density function and passing it to findPeaks() and findValleys() function of the quantmod R package^40^. The estimate of the intensity value of the valley was chosen as a threshold of minimal labelling signal and events below this value were discarded. (iv) Gating on cell size. Cells of homogeneous size were dynamically gated by pooling cells over all samples, defining a perimeter containing 75% of events in the FSC-A, SSC-A plane based on a density kernel and applying this gate to all samples. (v) Removal of doublets. As for cell-size, a gate that contained 80% of remaining events of all samples was then defined in the FSC-A, FSC-W plane using a density kernel function, and applied to all samples. (vi) Gating of cells in G1 phase of the cell-cycle. For each sample, a threshold of DAPI intensity was set at a value that minimized the derivative of the density function. Cells with intensity below this threshold were considered to be in the G1 peak of intensity and were kept for further analysis. (vii) Removal of samples with ≤ 1,000 events. At the end of these steps, about 10,000 cells remained on average per sample. (viii) Correction of fluorescent values for cell size. For each sample, a linear model of the form y ~ log(FSC.A) + log(SSC.A) was fitted using the MASS::rlm() function^41^ where y was the log intensity of fluorescence. Data of each cell *i* was then transformed as y_i_ = mean(y) + e_i_, where e_i_ was the *i*-th residual of the model. (ix) Variation between replicates was reduced by grouping all experiments of the same cell line and antibody labelling and applying on this group the flowStats::warpSet() function with default arguments. This function performed re-alignments according to high densities areas. (x) We observed that some samples displayed outlying distributions as compared to their replicated counterparts. To detect these cases in an unbiased way, we estimated all pairwise dissimilarities between replicates (same cell line and antibody staining) using the Kolmogorov-Smirnov statistics computed by the stats::ks.test() function. For each sample belonging to a group of at least three replicates, its dissimilarities to replicate counterparts were averaged and considered as a score K of reproducibility. Samples with K greater than the 95-th percentile of all K values were discarded.

### Analysis of flow cytometry data: traits describing cell-cell variability

Following data pre-processing, cell-to-cell variability within each sample was quantified by the coefficient of variation (CV = sd/mean) of the relevant fluorescent values. To account for sample-to-sample differences in mean expression levels, we also conditioned CV values on mean by computing the residuals of a non-parametric loess regression of CV ~ mean using the stats::loess() function. For CD23 which displayed bimodality, we fitted a 2 components gaussian mixture model (2GMM) on expression levels using the Mclust function from package mclust^42^ without constraint on parameters. This generated 5 parameters that fully described the distribution observed in each sample: mean and variance of the first component (μ_1_ and σ^2^_1_), mean and variance of the second component (μ_2_ and σ^2^_2_), and the proportion of cells (marginal weight) of the first component.

### Genetic linkage: genotypes dataset

The genotypes of 1000Genome individuals were downloaded from ftp://ftp.1000genomes.ebi.ac.uk/vol1/ftp/release/20130502/ on 13th February 2017. There were 40 individuals where genotyping was at phase 3 (NA19098, NA19099, NA19107, NA19108, NA19141, NA19204, NA19238, NA19239, NA18486, NA18488, NA18489, NA18498, NA18499, NA18501, NA18502, NA18504, NA18505, NA18507, NA18508, NA18516, NA18517, NA18519, NA18520, NA18522, NA18523, NA18853, NA18856, NA18858, NA18861, NA18867, NA18868, NA18870, NA18871, NA18873, NA18874, NA18912, NA18916, NA18917, NA18933, NA18934) and included phased genotypes (one file per chromosome of the hg19 genome release of February 2009, GRCh37 assembly). For 8 additional individuals (NA19140, NA19203, NA18487, NA18852, NA18855, NA18859, NA18862, NA18913), genotypes were unphased and obtained from ./supporting/hd_genotype_chip/ in the form of a single file with all chromosomes. Genotypes of 2 individuals were not found on the 1000Genome project server. Annotations of individuals (kinship and sexe) were obtained from file: ftp://ftp.1000genomes.ebi.ac.uk/vol1/ftp/release/20130502/integrated_call_samples_v2.20130502.ALL.ped. We used command lines G1-G4 of Supplementary Table S5 to extract genotypic data corresponding to individuals of our study. We selected variants located on a chromosomic region centered on the transcription start site (TSS) of each gene of interest. positions of these TSS were obtained from http://genome.ucsc.edu/cgi-bin/hgTables downloaded on 22-nd February 2017, using the ‘txStart’ field for genes CD55 and CD86 oriented in the forward direction, and the ‘txEnd’ field for genes CD23 and CD63 oriented in the reverse direction. Variants located within 2 Mb of the TSS were extracted with command line G5 of Supplementary Table S5. This produced 2 VCF files per gene, which contained the data of either 40 or 48 individuals, in a region of 4 Mb. These were converted to MAP and PED files using command G6 (Supplementary Table S5). Duplicated entries were removed by commands G7-G9 (Supplementary Table S5). Family ID and sex ID (1 for man and 2 for woman) were introduced in the ped file at the 1-st and 5-th column, respectively. Variants with MAF ≥ 0.05 were kept and variants that failed the Hardy-Weinberg test^43^ at a significance threshold of 0.001 were excluded using commands G10-11 (Supplementary Table S5).

### Genetic linkage: association test

We searched for genetic linkage between expression traits and markers, using either the 48 individuals with unphased genotypes or only the subset of 40 individuals with phased genotypes (see above). Expression traits for CD23, CD55, CD63 and CD86 were mean, CV and CV | mean; and for CD23 we also considered the five traits of the GMM model describing bimodal distributions (p_1_, μ_1_, μ_2_, σ_1_, σ_2_). We used PLINK (v1.9)^44^ to perform association tests, using either a purely additive model^45^ (command L1 of Supplementary Table S5) or a model that included possible dominance (command L2 of Supplementary Table S5). The model with dominance did not provide additional associations for variability and dispersion traits and all results of Table 1 were obtained with the additive model. Family-wise error rate across all variants was empirically estimated from 10,000 permutations and we retained association where this rate was lower than 0.05 (Table 1).

## ACKNOWLEDGEMENTS

We thank Mirko Francesconi and Olivier Gandrillon for critical reading of the manuscript, Thierry Defrance, Evelyne Manet, Jacqueline Marvel for discussions, Laura Presti and Celia Joseph for technical contribution, Véronique Barateau and SFR Biosciences Gerland-Lyon Sud (UMS344/US8) for access to flow cytometers and technical assistance, the 1000 Genome Project Consortium for access to cell lines and genotypes, the Pôle Scientifique de Modélisation Numérique for computing resources, BioSyL Federation and Ecofect Labex for inspiring scientific events, developers of IgBLAST, PLINK, R, git, bioconductor and Ubuntu for their software. This work was supported by the European Research Council under the European Union’s Seventh Framework Programme FP7/2007-2013 Grant Agreement n°281359 (GY) and EMBO long-term fellowship ALTF 691-2014 (OS).

## AUTHORS CONTRIBUTION

Performed experiments: G.T., O.S., H.G.

Developed analysis tools: C.B., S.J., G.Y.

Analysed the data and interpreted results: G.T., O.S., C.B., G.Y.

Conceived and supervised the study: G.Y.

Designed the study: O.S. and G.Y.

Wrote the paper: G.Y.

