## Supplemental Figures and Tables for "Cell-to-cell expression dispersion of B-cell surface proteins displays genetic variation among humans"

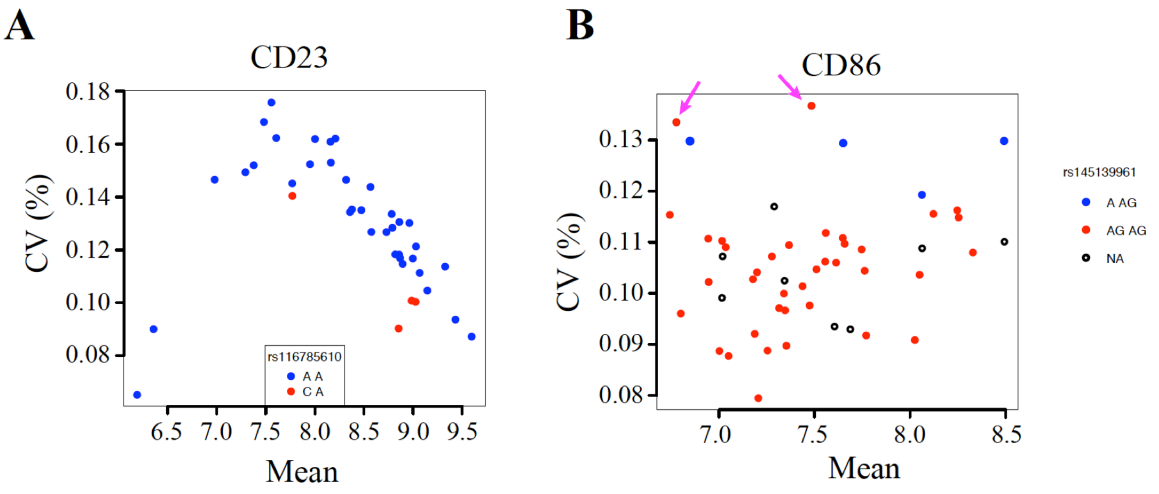


**Supplementary Figure S1. A)** Dot plot of CV vs. mean expression of CD23 in 40 LCLs, colored according to their genotype at SNP rs116785610. Association with expression dispersion was significant at $p$ = $10^{-4}$ nominally and at $p$ = 0.043 when accounting for the number of SNPs tested. Note that linkage was searched for eight CD23 expression traits (mean, variability, dispersion and 5 traits describing bimodality, see methods). **B**) Dot plot of CV vs. mean expression of CD86, colored according to genotype at SNP rs145139961. NA: genotype not available. Arrows: Genotypes determined by PCR, rejecting association initially detected (Table 1).

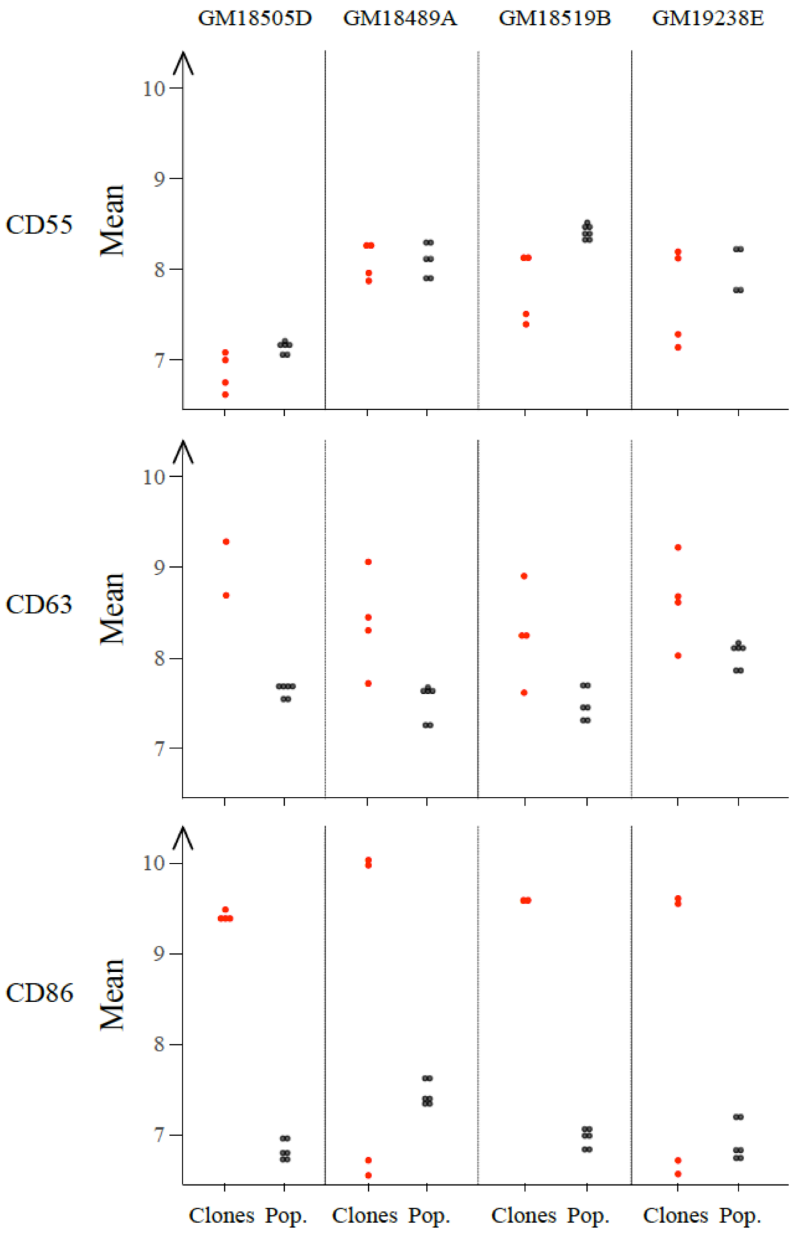


**Supplementary Figure S2. Technical shift between two series of acquisitions.** The survey of 50 LCLs (Pop., black dots) and the analysis of subclones (Clones, red dots) were performed in two distinct series of acquisitions, about 6 months apart. Four cell lines (indicated on top) were processed in both analyses. For CD63 and CD86, we observed a shift in mean values between the Pop. series and most samples of the Clones series, preventing reliable comparisons between the two series.

**Supplementary Table S1: CDR3 genotypes obtained from clonality PCR.** Numbers x/y are the percentages of counts corresponding to the CDR3 sequences indicated in columns, for samples of cell lines indicated in rows; x and y being the percentage in the first and second replicate sample of the cell line, respectively. Only sequences represented by at least 20% of the counts, based on at least 20 counts, are represented. (-) not detected in this sample; (i) sequence TSGNTGWYSDYWGQG corresponding to GM18505D cell line and subclones seems to have contaminated other samples; (ii) of a total of less than 100 counts; (*) of a total of more than 130,000 counts.

**1A**

| | TSGNTGWYSDYWGQG (i) | **Conclusion** |
|---|---|---|
| GM18505D | 85.7(*)/85.8(*) | monoclonal |
| 5D-4F9 | 94.7(*)/94.9(*) | monoclonal |
| 5D-5D10 | 95.5(*)/95.2(*) | monoclonal |

**1B**

| | AKHYDYWGQG | **Conclusion** |
|---|---|---|
| GM18486A | 57.7(*)/52.5(*) | monoclonal (multiple other sequences were detected but did not reach 20% of representativity) |
| 6A-1C10 | 95.8(*) | monoclonal |
| 6A-2E9 | 95.3(*)/96.0(*) | monoclonal |
| 6A-4F9 | 96.0(*)/96.2(*) | monoclonal |
| 6A-5C3 | 97.0(*)/96.1(*) | monoclonal |

**1C**

| | ARGVMYDSGLISGWGQG | ARVGANNWYENDYWGQG | ARDLVAAAGTLGYWGQG | AKHYDYWGQG | TSGNTGWYSDYWGQG (i) | OTHERS | **Conclusion** |
|---|---|---|---|---|---|---|---|
| GM18519B | 39.1/47 | | | | | 61.9/53 | polyclonal |
| 9B-3G7 | | 80.5/95.6 | | | | | monoclonal |
| 9B-5B4 | | 92.9/94.2 | | | | | monoclonal |
| 9B-5D4 | | 84.8/92.4 | | | | | monoclonal |
| 9B-5E5 | | 84.7/94.1 | | | | | monoclonal |
| 9B-2F4 | -/44.0(ii) | 94.0/ - | | | | | monoclonal |
| 9B-4C3 | | 93.9/81.5 | | | | | monoclonal |
| 9B-4G9 | | 92.4/84.8 | | | | | monoclonal |
| 9B-5G5 | | 94.0/86.3 | | | | | monoclonal |
| 9B-1D10 | | | 22.3 | 56.9 | | | polyclonal |
| 9B-2F10 | | 91.9/ 66.5 | | | -/26.0 | | monoclonal (i) |
| 9B-2B5 | | 91.5/55.5 | | | -/39.3 | | monoclonal (i) |

**1D**

| | ARHHNYYDNSPEDWFDSWGHG | ARVGANNWYENDYWGQG | **Conclusion** |
|---|---|---|---|
| GM19239C | 44.1/42.9(*) | | unclear (one major sequence but not found in sequences from subclones) |
| 9C-2F5 | | 54.8(ii) / - | unclear (low coverage) |
| 9C-6G5 | | 50(ii) | unclear (low coverage) |

**1E**

| | VRDGGGWWYLYWGQG | AREEYSGSYYGYFQHWGQG | SRENDYSRSPEYWGQG | **Conclusion** |
|---|---|---|---|---|
| GM18489A | 44.8/38.8 | 23.8/25.6 | | polyclonal |
| 9A-3H10 | | | 92.3(*)/93.3(*) | monoclonal |

**1F**

| | ARDLVAAAGTLGYWGQG | ARDPAAGMQYFQHWGQG | AKVSAEFSTNWTFDFWGQG | AKAAMYYDSSGYMIFGGRWGQG | TSGNTGWYSDYWGQG (i) | AKHYDYWGQG | ARHHNYYDNSPEDWFDSWGHG | PAQYYFGYWGQGTLVTV | **Conclusion** |
|---|---|---|---|---|---|---|---|---|---|
| GM19238E | 41.5/ 42.2 | 31.0/ 30.4 | | | | | | | polyclonal |
| 8E-3F7 | | | 89.4/ 90.3 | | | | | | monoclonal |
| 8E-8E4 | | | | 83.3/34.2 | - / 59.1 | | | | monoclonal (i) |
| 8E-2G6 | | | | 89.6 | | | | | monoclonal |
| 8E-4G4 | | | | 95.9/95.7 | | | | | monoclonal |
| 8E-6C10 | | | | | 31.7 (ii)/ 93.1 (ii) | 26.8/ - | 26.8/ - | | polyclonal |
| 8E-7E10 | | | | | 54.0 (ii) / 41.6 (ii) | | | 27.0/ 37.7 | unclear (i) |
| 8E-1G6 | | | | 25.2 | 71.2 | | | | unclear (i) |

**Supplementary Table S2: Primers used for CDR3 amplification and sequencing**

| ID | 5'-3' sequence |
|---|---|
| 1K47 | TCGTCGGCAGCGTCAGATGTGTATAAGAGACAGTCTCCAAATGAACAGCCTGAGAGCC |
| 1K48 | TCGTCGGCAGCGTCAGATGTGTATAAGAGACAGGAGCTCTGTGACCGCCGCGGACACG |
| 1K49 | GTCTCGTGGGCTCGGAGATGTGTATAAGAGACAGACCTGAGGAGACGGTGACC |
| 1K71 | TCGTCGGCAGCGTCAGATGTGTATAAGAGACAGTGGAGCTGAGCAGCCTGAGATCTGA |
| 1K72 | TCGTCGGCAGCGTCAGATGTGTATAAGAGACAGCAATGACCAACATGGACCCTGTGGA |
| 1K73 | TCGTCGGCAGCGTCAGATGTGTATAAGAGACAGCAGCACCGCCTACCTGCAGTGGAGC |
| 1K74 | TCGTCGGCAGCGTCAGATGTGTATAAGAGACAGGTTCTCCCTGCAGCTGAACTCTGTG |
| 1K75 | TCGTCGGCAGCGTCAGATGTGTATAAGAGACAGCAGCACGGCATATCTGCAGATCAG |

**Supplementary Table S3. IgBLAST command lines used to extract CDR3 sequences.**

| ID | Command | Description |
|---|---|---|
| C1 | makeblastdb -title human_gl_V -<br>parse_seqids -dbtype nucl -in<br>'"human_gl_VH_IgGermline.fasta"<br>"human_gl_VK_IgGermline.fasta"<br>"human_gl_VL_IgGermline.fasta"'<br>-input_type fasta -out<br>human_gl_V -hash_index | Create BLAST database of human V segments |
| C2 | makeblastdb -title human_gl_D -<br>parse_seqids -dbtype nucl -in<br>'"human_gl_DH_IgGermline.fasta"'<br>-input_type fasta -out<br>human_gl_D -hash_index | Create BLAST database of human D segments |
| C3 | makeblastdb -title human_gl_J -<br>parse_seqids -dbtype nucl -in<br>'"human_gl_JH_IgGermline.fasta"<br>"human_gl_JK_IgGermline.fasta"<br>"human_gl_JL_IgGermline.fasta"'<br>-input_type fasta -out<br>human_gl_J -hash_index | Create BLAST database of human J segments |
| C4 | igblastn -query assembled.fasta<br>-germline_db_V human_gl_V -<br>germline_db_D human_gl_D -<br>germline_db_J human_gl_J -<br>auxiliary_data<br>./optional_file/human_gl.aux -<br>organism human -domain_system<br>imgt -outfmt 3 -show_translation<br>-num_alignments_V 1 -<br>num_alignments_D 1 -<br>num_alignments_J 1 | Assign every read to one gene segment following the IMGT ontology (http://www.imgt.org) |

**Supplementary Table S4: List of antibodies used in this study**

| Human antigen | Host | Fluorescent Conjugate | Provider | Catalog Nb | Other |
|---|---|---|---|---|---|
| CD2 | mouse | APC | BioLegend | 300213 | clone RPA-2.10, IgG1, κ |
| CD5 | mouse | PE/Cy7 | BioLegend | 300622 | clone UCHT2, IgG1, κ |
| CD9 | mouse | FITC | BioLegend | 312104 | clone Hl9a, IgG1, κ |
| CD19 | mouse | Alexa647 | Invitrogen Molecular Probes | MHCD1921 | clone SJ25-C1, IgG1 |
| CD20 | mouse | FITC | BioLegend | 302304 | clone 2H7, IgG2b, κ |
| CD22 | mouse | APC | BioLegend | 302510 | clone HlB22, IgG1, κ |
| CD23 | mouse | FITC | BD Pharmingen | 561146 | clone ML233, IgG1, κ |
| CD23 | mouse | APC | BD BioSciences | N/A | clone ML233 at 100 μg/ml. Dilution used: 3.75/1000 |
| CD37 | mouse | FITC | BioLegend | 356303 | clone M-B371, IgG1, κ |
| CD38 | mouse | PE/Cy7 | BioLegend | 303515 | clone HlT2, IgG1, κ |
| CD40 | mouse | PE/Cy7 | BioLegend | 334321 | clone 5C3, IgG1, κ |
| CD46 | mouse | FITC | BD Pharmingen | 555949 | clone E4.3, IgG2a, κ |
| CD53 | mouse | PE | BioLegend | 325406 | clone Hl29, IgG1, κ |
| CD55 | mouse | APC | BioLegend | 311311 | JS-11, IgG1, κ at 100 μg/ml. Dilution used: 3.75/1000 |
| CD59 | mouse | FITC | LifeTech/Invitrogen Molecular Probes | MHCD5901 | clone MEM-43, IgG2a |
| CD63 | mouse | PE/Cy7 | BioLegend | 353009 | clone H5C6, |

| | | | | | |
|---|---|---|---|---|---|
| | | | | | IgG1, κ at 200 µg/ml. Dilution used: 4/1000 |
| CD79b | mouse | PE | BD Pharmingen | 557931 | clone 3A2-2E7, IgG1, κ |
| CD80 | mouse | Alexa647 | BioLegend | 305215 | clone 2D10, IgG1, κ |
| CD86 | mouse | PE | BioLegend | 305405 | IT2.2, IgG2b, κ at 100 µg/ml. Dilution used: 3.75/1000 |

**Supplementary Table S5. Command lines used for linkage analysis.**

<table>
<tr><th>ID</th><th>Command</th><th>Description</th></tr>
<tr><td>G1</td><td>tabix -p vcf</td><td>indexing VCF files (tabix v0.2.6)[1]</td></tr>
<tr><td>G2</td><td>perl vcf-subset -c „40 LCLs IDs“ variants_chr.vcf.gz | bgzip -c > 40LCLs_variants_chr.vcf.gz</td><td>subsetting VCF files for the set of 40 individuals with phased genotypes (vcftools v0.1.13)[2]</td></tr>
<tr><td>G3</td><td>perl vcf-subset -c „8 LCLs IDs“ variants_chrall.vcf.gz | bgzip -c > 8LCLs_variants_chrall.vcf.gz</td><td>subsetting VCF files for the set of 8 individuals with unphased genotypes (vcftools v0.1.13)[2].</td></tr>
<tr><td>G4</td><td>perl vcf-merge 40LCLs_variants_chr.vcf.gz 8LCLs_variants_chrall.vcf.gz | bgzip -c > 48LCLs_variants_chr.vcf.gz</td><td>Merging variants that are genotyped and uniquely identified in all 48 individuals (vcftools 0.1.13).</td></tr>
<tr><td>G5</td><td>tabix -hp vcf chr.vcf.gz chr:begin-end | .bgzip -c > deltaTSS_chr.vcf.gz</td><td>Select variants in regions flanking genes TSS.</td></tr>
<tr><td>G6</td><td>vcftools --gzvcf deltaTSS_chr.vcf.gz –plink</td><td>Convert VCF files into MAP and PED files</td></tr>
<tr><td>G7</td><td>plink --file deltaTSS_chr --list-duplicate-vars</td><td>removal of variants present in duplicates. (PLINK v1.9)[3]</td></tr>
<tr><td>G8</td><td>plink –file deltaTSS_chr –exclude „IDs to remove“ --make-bed –out clean</td><td>further removal of variants with non-unique IDs (duplicated IDs in second colmun of MAP file)</td></tr>
<tr><td>G9</td><td>plink –bfile clean --recode –tab –out clean_nodup</td><td>build final dataset</td></tr>
<tr><td>G10</td><td>plink --bfile clean_nodup --maf 0.05 --hwe 0.001 --make-bed --</td><td>Exclude variants with MAF < 0.05 or deviating from Hardy-Weinberg.</td></tr>
</table>

| | | |
|---|---|---|
| | `out clean_nodup_filt` | |
| G11 | `plink --bfile clean_nodup_filt --recode --tab --out clean_nodup_filt_tab` | Re-encode data for PLINK after cleanup |
| L1 | `plink –bfile clean_nodup_filt_tab --assoc qt-means --mperm 10000 –out lm` | Linkage test without dominance. |
| L2 | `plink –bfile clean_nodup_filt_tab --linear genotypic --mperm 10000 –out lm` | Linkage test allowing dominance. |
